# Measuring Sequences of Representations with Temporally Delayed Linear Modelling

**DOI:** 10.1101/2020.04.30.066407

**Authors:** Yunzhe Liu, Raymond J Dolan, Hector Luis Penagos-Vargas, Zeb Kurth-Nelson, Timothy Behrens

## Abstract

There are rich structures in off-task neural activity. For example, task related neural codes are thought to be reactivated in a systematic way during rest. This reactivation is hypothesised to reflect a fundamental computation that supports a variety of cognitive functions. Here, we introduce an analysis toolkit (TDLM) for analysing this activity. TDLM combines nonlinear classification and linear temporal modelling to testing for statistical regularities in sequences of neural representations. It is developed using non-invasive neuroimaging data and is designed to take care of confounds and maximize sequence detection ability. The method can be extended to rodent electrophysiological recordings. We outline how TDLM can successfully reveal human replay during rest, based upon non-invasive magnetoencephalography (MEG) measurements, with strong parallels to rodent hippocampal replay. TDLM can therefore advance our understanding of sequential computation and promote a richer convergence between animal and human neuroscience research.

## INTRODUCTION

Human neuroscience has made remarkable progress in detailing the relationship between the representations of different stimuli during task performance ^1,2^. At the same time, it is increasingly clear that resting, off-task, brain activity is structurally rich and is important for understanding the neural underpinnings of cognition ^3^. However, unlike the case for task-based activity, little attention has been given to techniques that can measure representational content or structure of this resting activity. Here, we introduce TDLM (temporal delayed linear modelling) as an analysis framework, based on linear modelling, that can characterize temporal structure of internally generated neural representations.

TDLM enables a detailed examination of sequential patterns in neural code reactivation that are not tied to task events. Our approach is inspired by evidence from the rodent literature of rich temporal structure in representational content of offline brain activity. Here a seminal finding in rodent electrophysiological research is “hippocampal replay” ^4–6^. During rest and quiet wakefulness, place cells in the hippocampus (that signal self-location during periods activity) spontaneously recapitulate old, and explore new, trajectories through an environment ^4,5^. These internally generated sequences are hypothesized to reflect a fundamental feature of neural computation across tasks ^7–10^.

Applying TDLM on non-invasive neuroimaging data we, and others, have shown it is possible to measure spontaneous sequences of neural representations during rest in humans ^11,12^. The results resemble key characters found in rodent hippocampal replay and inform key computational principles of human cognition ^12^.

In the following sections, we introduce the logic and mechanics of TDLM in detail. We first compare performance of alternative algorithms on synthetic data where the ground truth is known (see detailed description of the synthetic data and simulation code in Supplementary Note 1). Subsequently, we apply the method to real neural data, both human magnetoencephalography (MEG) ^11,12^ and rodent hippocampal electrophysiological recordings (Supplementary Note 2). In relation to the latter, we show TDLM successfully reproduces key findings, including the presence of theta sequences ^13^.

TDLM is a general, and flexible, tool for measuring neural sequences. It facilitates crossspecies investigations by linking large-scale measurements in humans to cellular measurements in non-human species. We outline its promise for revealing abstract cognitive processes that extend beyond sensory representation, potentially opened doors for new avenues of research in cognitive science. All code and facilities will be available at https://github.com/yunzheliu/TDLM.

## RESULTS

### TDLM

#### Overview of TDLM

Our primary goal is to test for temporal structure in neural activity. To achieve this, we would like ideally a method which (1) uncovers regularity in the reactivation of neural activity, (2) tests whether this regularity conforms to a hypothesized structure. Here the structure between neural representation is expressed as their sequential reactivation in time, i.e., sequence. In what follows, we will use the terms “temporal structure” and “sequence” interchangeably.

The starting point of TDLM is a set of *n* time series, each corresponding to a decoded neural representation of a variable of interest. These time series could themselves be obtained in several ways, described in detail in a later section (“Getting the states”). The aim of TDLM is to identify task-related regularities in sequences of these representations off-task.

Consider, for example, a task in which participants have been trained such that *n*=4 distinct sensory cues (A, B, C, and D) appear in a consistent order (*A → B → C → D*) (Fig 1a). If we are interested in replay of this sequence during subsequent resting periods, we might want to ask statistical questions of the following form: “Does the existence of a neural representation of A, at time T in the rest period, predict the occurrence of a representation of B at time T+Δ*t*”, and similarly for *B → C* and *C → D*.

**Fig. 1.**
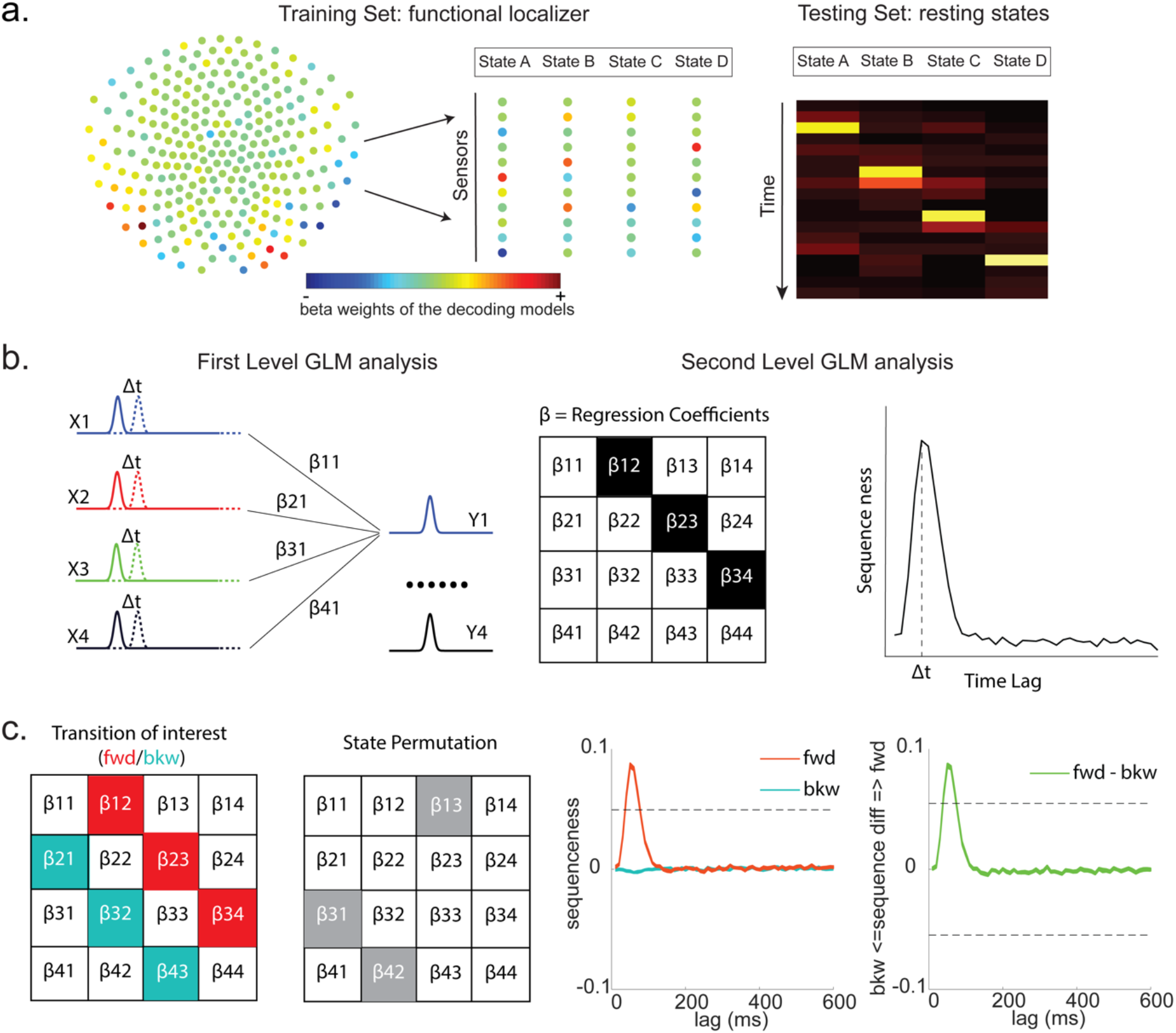
TDLM is a linear generative model that assumes that state time course can be predicted by other lagged state time courses. **a,** State definition: The first step of TDLM is decoding, to establish the mapping between multivariate neural patterns and labelled states through supervised learning. A separate decoding model, e.g. regularised logistic regression was trained to recognize each state (left) vs. other states and null data. Decoding models (consisting of a set of weights over sensors) were then tested on unlabelled testing data (e.g., resting state) to generate a time*state decoding matrix (right). Examples of forward sequential state reactivations in simulated data (right). **b,** The second step of TDLM is to quantify the temporal structure of the decoded states using a two-level GLM approach. In the first level GLM result in a state*state regression coefficient matrix at each time lag. In the second level GLM, this coefficient matrix is projected onto the hypothesized state transition matrix (in red), to give a single measure of sequenceness as a function of time-lag. **c,** The statistical significance was tested using a nonparametric state permutation test by randomly shuffling the transition matrix of interest (in grey). The statistical significance threshold is defined as the 95^th^ percentile of all shuffles across all time lags for raw forward (in red) and backward (in blue) sequence, denoted as the dashed line. In addition, we define a summary statistic – sequence difference (D), which is the subtraction of forward and backward sequence at each time lag (in green). Positive value means favouring forward vs. backward, and vice versa. The permutation threshold for sequenceness is defined as the 95^th^ percentile of the maximum absolute value of the subtraction of all shuffles across all time lags.

In TDLM we ask such questions using a two-step process. First, for each of the *n*^2^ possible pairs of variables *X_i_* and *X_j_*, we find the correlation between the *X_i_* time series and the Δ*t*-shifted *X_j_* time series. These *n*^2^ correlations comprise an empirical transition matrix, describing how likely each variable is to be succeeded at a lag of Δ*t* by each other variable (Fig. 1b, left panel). Second, we correlate this empirical transition matrix with a task-related transition matrix of interest (Fig. 1b, right panel). This produces a single number that characterizes the extent to which the neural data follow the transition matrix of interest, which we call ‘sequenceness’. Finally, we repeat this entire process for all Δ*t* > 0, yielding a measure of sequenceness at each possible lag between variables (Fig. 1c).

Note that, for now, this approach decomposes a sequence (such as *A → B → C → D*) into its constituent transitions and adds the evidence for each transition. It therefore does not require that the transitions themselves are sequential: *A → B* and *B → C* could occur at unrelated times, so long as the within-pair time lag was the same. In section “Multi-step sequences”, we address how to strengthen the inference by looking explicitly for longer sequences.

#### Constructing the empirical transition matrix

In order to find evidence for state-to-state transitions at some time lag Δ*t*, we could regress a time-lagged copy of one state, *X_j_*, onto another, *X_i_*:

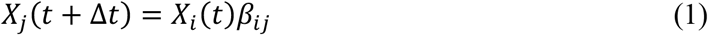

Instead, TDLM includes all states in the same regression model for important reasons, detailed in section “Moving to multiple linear regression”:

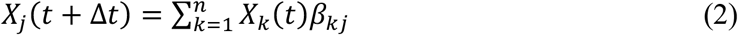

In this equation, the values of all states *X_k_* at time *t* are used in a single multilinear model to predict the value of the single state *X_j_* at time *t* + Δ*t*.

The regression described in Equation 2 is performed once for each *X_j_*, and these equations can be arranged in matrix form as follows:

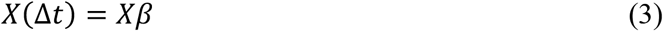

Each row of *X* is a timepoint, and each of the *n* columns is a state. *X(*Δ*t*) is the same matrix as *X*, but with the rows shifted forwards in time by Δ*t*. *β* is an *n* × *n* matrix of weights - which we call the *empirical transition matrix*. *β_ij_* is an estimate of the influence of *X_i_*(*t*) on *X_j_*(*t* + Δ*t*), over and above variance that can be explained by other states at the same time.

To obtain *β*, we invert Equation 3 by ordinary least squares regression.

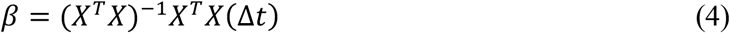

This inversion can be repeated for each possible time lag (Δ*t* = 1,2,3, …), resulting in a separate empirical transition matrix *β* at every time lag. We call this step the first level sequence analysis.

#### Testing the hypothesized transitions

The first level sequence analysis assesses evidence for all possible state-to-state transitions. The next step in TDLM is to test for the strength of a particular hypothesized sequence, specified as a transition matrix, *T_F_*. We therefore construct another GLM which relates *T_F_* to the empirical transition matrix *β*. We call this step the second level sequence analysis:

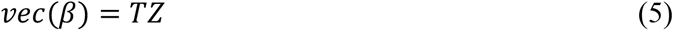

where *vec* denotes the vectorized form of a matrix, *T* is the design matrix. *T* has 4 columns, each of which is a vectorized transition matrix: *vec*(*T_auto_*), *vec*(*T_Const_*), *vec*(*T_F_*) and *vec*(*T_B_*). *T_F_* is the task-related transition matrix of interest, as described above. *T_B_* is *T_F_* transposed; that is, the same transitions in the backward direction. *T_Const_* is a constant vector that models away the average of all transitions, ensuring that any weight on *T_F_* and *T_B_* is specific to the hypothesized transitions. *T_auto_* models self-transitions to control for autocorrelation (equivalently, we could simply omit the diagonal elements from the regression). *Z* is the weights of the second level regression, which has four entries. Repeating the regression of Equation 5 at each time lag (Δt = 1,2, 3, …) results in four vectors, which we will call *Z_F_*, *Z_B_*, *Z*uto*, and *Z_Const_*. The values of *Z_F_* and *Z_B_* at each time lag are our estimates of forward and backward sequence strength (respectively) at that lag, along the transition matrix of interest (Fig. 1c).

In many cases, *Z_F_* and *Z_B_* will be the final outputs of a TDLM analysis. However, it may sometimes also be useful to consider the quantity:

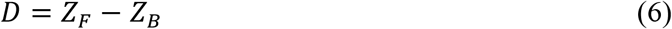

*D* contrasts forward and backward sequences to give a measure that is positive if sequences occur mainly in a forward direction and negative if sequences occur mainly in a backward direction. This may be advantageous if, for example, *Z_F_* and *Z_B_* are correlated across subjects (due to factors such as subject engagement and measurement sensitivity). In this case, *D* may have lower cross-subject variance than either *Z_F_* or *Z_B_*, as the subtraction removes common variance.

Finally, to test for statistical significance, TDLM relies on a nonparametric permutation-based method. The null distribution is constructed by randomly shuffling the identities of the *n* states and re-calculating the second level analysis for each shuffle. The first level analysis retains the veridical labels. This approach allows us to reject the null hypothesis that there is no relationship between the empirical transition matrix and the task-defined transition of interest. Note that there are many wrong ways to perform permutations, which permute factors that are not exchangeable under the null hypothesis and therefore lead to false positives. We will examine some of these later with simulations. In some cases, it may be desirable to test slightly different hypotheses by using a different set of permutations; this will also be discussed later.

If the time lag Δ*t* at which neural sequences exist is not known *a priori*, then we must correct for multiple comparisons over all tested lags. This can be achieved by using the maximum *Z_F_* across all tested lags as the test statistic. If we choose this test statistic, then any values of *Z_F_* exceeding the 95^th^ percentile of the null distribution can be treated as significant at *α* = 0.05.

### TDLM STEPS IN DETAIL

#### Getting the states

As described above, the input to TDLM is a set of time series of decoded neural representations, or states. Here we give three examples of specific state spaces that we have worked with using TDLM.

#### States as sensory stimuli

The simplest case, perhaps, is to define a state in terms of a neural representation of sensory stimuli, e.g., face, house. To obtain their neural representation, we present the stimuli in a randomized order at the start of a task, while whole-brain neural activity is recorded by non-invasive neuroimaging method, e.g., MEG or EEG. We then train a supervised decoding model to map the pattern of recorded neural activity to the presented image (Supplementary Fig. 1).

This could be any of the multitude of available decoding models. For simplicity we have used a logistic regression model throughout.

In MEG/EEG, neural activity is recorded by multiple sensor or channel arrays on the scalp. The sensor arrays record whole-brain neural activity at millisecond temporal resolution. To avoid potential selection bias (given the sequence is expressed in time), we choose to use the whole brain sensor activity at a single time point (i.e., spatial feature) as the training data fed into classifier training.

Ideally, we would like to select a time point where the neural activity can be most truthfully read out. This can be indexed as the time point that gives the peak decoding accuracy. If the state is defined by the sensory feature of stimuli, we can use a classical leave-one-out crossvalidation scheme to determine the ability of classifiers to generalise to unseen data of the same stimulus type (decoding accuracy) at each time point. This cross-validation scheme is asking whether the classifier trained on the sensory feature can be used to classify the unseen data of same stimuli (Fig. 2a, b).

**Fig. 2.**
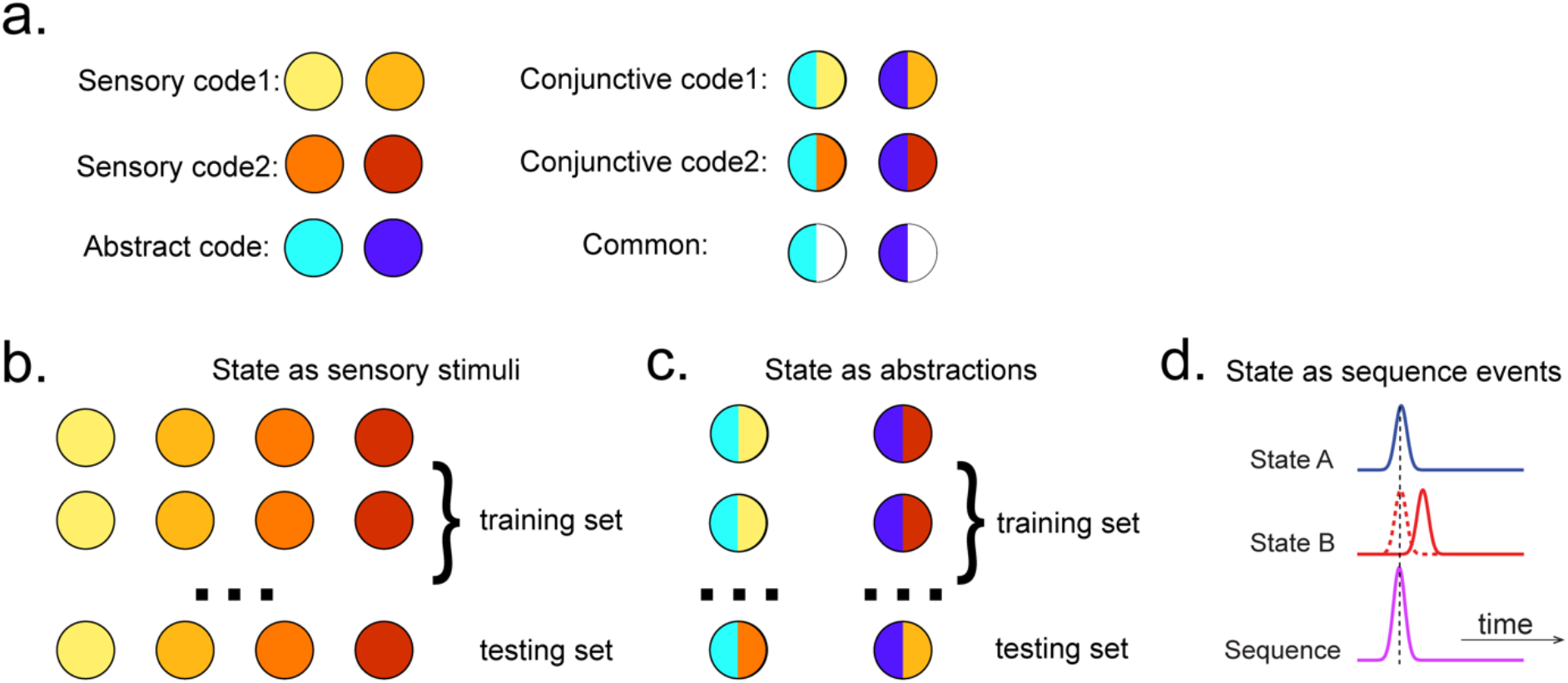
TDLM is capable of working on different state spaces. **a**, Assuming we have two abstract codes, each abstract code has two different sensory codes (left panel). The M/EEG data corresponding to each stimulus is a conjunctive representation of sensory and abstract codes (right panel). The abstract code can be operationalised as the common information in the conjunctive codes of two stimuli that share the same abstract representation. **b**, Training decoding models for stimulus information. The simplest state is defined by sensory stimuli. To determine the best time point for classifier training, we can use classical leave-one-out cross validation scheme on the stimuli-evoked neural activity. **c**, Training decoding models for abstracted information. The state can also be defined as the abstractions. To extract this information only, we need to avoid sensory information. We can train the classifier on the neural activity evoked by one stimulus and tested on the other sharing the same abstract representation. If the neural activity contains both the sensory and abstract code, then the only information can generalize is the common abstract code. **d**, The state can also be defined as the sequence event itself.

#### States as abstractions

As well as sequences of sensory representations, it is possible to search for replay of more abstract neural representations, within the constraint that we can build a decoder for them. Such abstractions might be associated with the presented image (e.g., mammal vs fish), in which case analysis can proceed as above by swapping categories for images.

A more subtle example, however, is where the abstraction has to do with the sequence or graph itself. For example, one representation of interest might be whatever is common at a particular location in space but invariant to what sensory stimuli are present at that location ^14^ A related type of abstraction corresponds to the position of an item in a sequence, invariant to which actual item is in that position ^12,15^.

We need to exercise care when setting up cross-validation schemes for training “abstract” classifiers, because we don’t want the “abstract” classifier to capitalize on common sensory features. Otherwise, we might report false positive sequences of abstract codes when in fact there is only sequence for sensory information (Supplementary Fig. 2). This can happen if we train and test on the same sensory (as well as abstract) object. In other words, we need to ensure that there is no one-to one mapping between sensory and abstract code. To do so, we need more than one sensory exemplar of each abstract state.

If we have exemplars of *N (N >* 1) different sensory images for each abstract state, then training can proceed in the following way. For example, the training set for the “2” decoder comprises *N − 1* types of sensory images at position 2, leaving out all instances of one single type sensory example for cross-validation. Therefore, an above chance classification must rely on features that are shared between the N-1 sensory images and the one left-out sensory image, which is the abstract code. If there are just 2 stimuli per abstraction, we can train on one stimulus, and test on the other (and vice versa), selecting the time point that does best in this “cross-validation”. This scheme therefore searches for representations that generalise over at least two stimuli that embody the same abstract meaning (Fig. 2c).

#### States as sequence events

TDLM can also be used iteratively to ask question about the ordering of different types of replay events (Fig. 2d). This can lead to powerful inferences about the temporal organisation of replay, such as “Rapid replay of sensory representations is embedded within a lower frequency rhythm”. This more sophisticated use of TDLM merits its own consideration and is discussed below under “Sequences of sequences”.

#### Controlling confounds and maximising sensitivity in sequence detection

Here, we motivate the key features of TDLM.

#### Temporal correlations

In standard linear methods, unmodelled temporal autocorrelation can inflate statistical scores. Techniques such as auto-regressive noise modelling are commonplace to mitigate these effects ^16,17^ However, autocorrelation is a particular burden for analysis of sequences, where it interacts with correlations between the decoded neural variables.

To see this, consider a situation where we are testing for the sequence *X_i_ → X_j_*. TDLM is interested in the correlation between *X_i_* and lagged *X_j_* (see Equation 1). But if the *X_i_* and *X_j_* time series contain autocorrelation and are also correlated with one another, then *X_i_*(*t*) will necessarily be correlated with *X_j_*(*t* + Δ*t*). Hence, the analysis will spuriously report sequences.

Correlations between states are commonplace. Consider representations of visual stimuli decoded from neuroimaging data. If these states are decoded using an *n*-way classifier (forcing exactly one state to be decoded at each moment), then the *n* states will be anti-correlated by construction. On the other hand, if the states are each classified against a null state corresponding to the absence of stimuli, then the *n* states will typically be positively correlated with one another.

Notably, in our case, because these autocorrelations are identical between forward and backward sequences, one approach for removing them is to compute the difference measure described above (*D* = *Z_F_* − *Z_B_*). This approach that works well was suggested in Kurth-Nelson, et al. ^11^. However, a downside it that it prevents us from measuring forward and backward sequences independently. The remainder of this section considers alternative approaches that can allow independent measurement of forward and backward sequences.

#### Moving to multiple linear regression

The spurious correlations above are induced because *X_j_*(*t*) mediates a linear relationship between *X_i_*(*t*) and *X_j_*(*t* + Δ*t*). Hence, if we knew *X_j_*(*t*), we could solve the problem by simply controlling for it in the linear regression, as in Granger Causality ^18^:

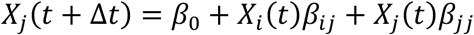

Unfortunately, however, we do not have access to the ground truth of *X* - since these variables have been decoded noisily from brain activity. Any error in *X_j_*(*t*) but not *X_i_*(*t*) will mean that the control for autocorrelation will be imperfect, leading to spurious weight on *β_ij_*, and therefore spurious inference of sequences.

This problem cannot be solved without a perfect estimate of *X*, but it can be systematically reduced until negligible. It turns out the necessary strategy is simple. We do not know ground truth *X_j_*(*t*), but what if we knew a subspace that included estimated *X_j_*(*t*)? If we controlled for that whole subspace, we would again be safe. We can get closer and closer to this by including further co-regressors that are themselves correlated with estimated *X_j_*(*t*) with different errors from ground truth *X_j_*(*t*). The most straightforward approach is to include the other states of *X*(*t*), each of which has different errors, leading to the multiple linear regression of Equation 2.

Figure 3a shows this method applied to the same simulated data whose correlation structure induces false positives in the simple linear regression of Equation 1. The multiple regression accounts for the correlation structure of the data and allows correct inferences to be made. Unlike the simple subtraction method proposed above (Fig. 3a, left panel), the multiple regression permits separate inference on forwards and backwards sequences.

**Fig. 3.**
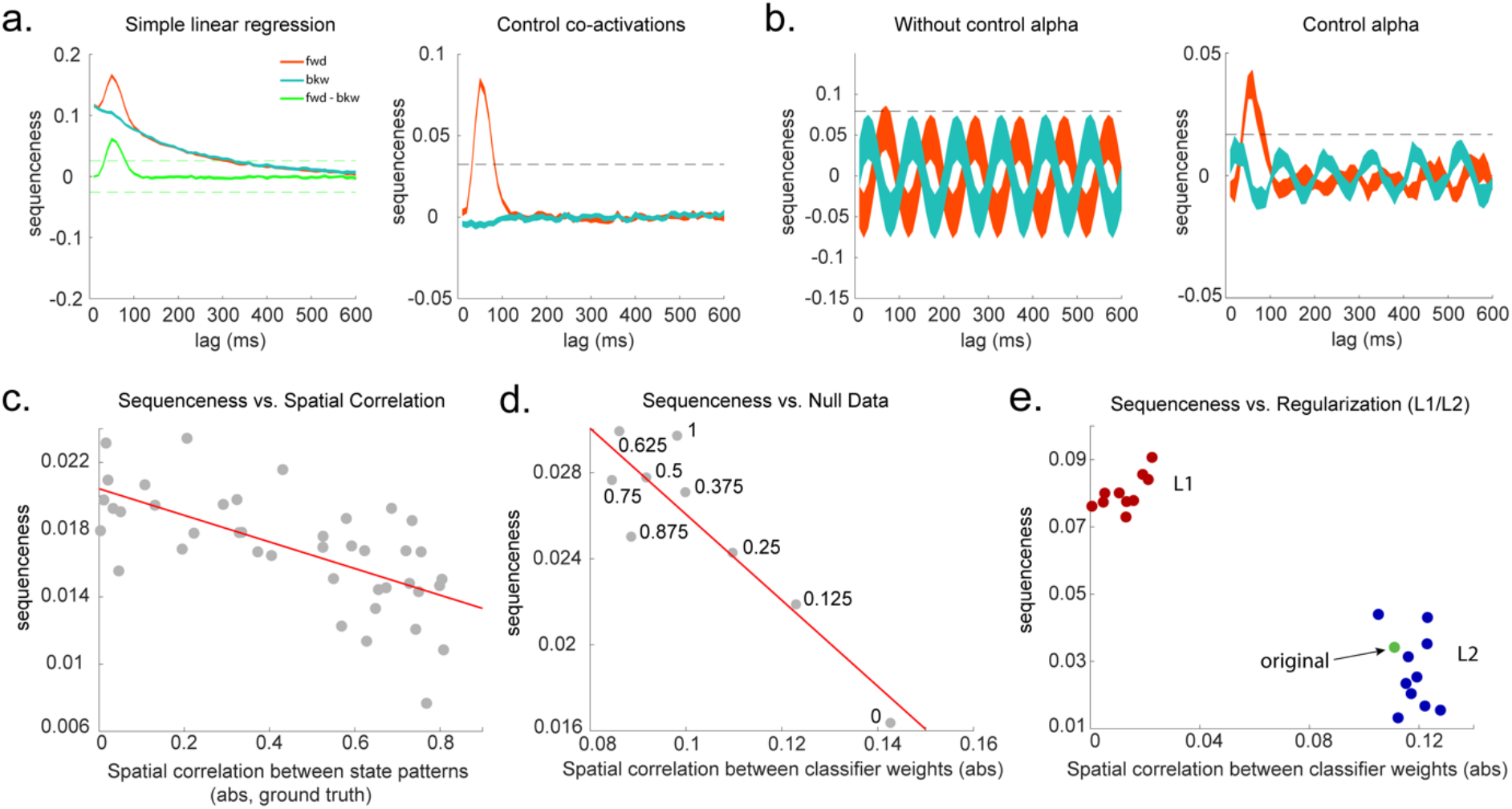
TDLM builds a linear model to test for sequential structure in state reactivations by controlling for temporal and spatial correlations. **a,** Simple linear regression or cross-correlation approach relies on the asymmetry of forward and backward transitions; therefore, subtraction is necessary (left panel). TDLM instead relies on multiple linear regression. TDLM can assess forward and backward transitions separately (right panel). **b,** Background alpha oscillations, as seen during rest periods, can reduce sensitivity of sequence detection (left panel), controlling alpha in TDLM helps recover the true signal (right panel). **c,** The spatial correlation between the sensor weights of decoders for each state reduces the sensitivity of sequence detection. This suggests reducing overlapping patterns between states are important for sequence detection. **d,** Adding null data to the training set can help increase the sensitivity of sequence detection by reducing the spatial correlations of the trained classifier weights. Here the number indicates the ratio between null data and task data. “1” means the same amount of null data and the task data. “0” means no null data is added for training. **e,** L1 regularization helps sequence detection by reducing spatial correlations (all red dots are L1 regularization with a varying parameter value), while L2 regularization does not help sequenceness (all blue dots are L2 regularization with a varying parameter value) as it does not reduce spatial correlations of the trained classifiers compared to the classifier trained without any regularization (green point).

#### Oscillations and long timescale autocorrelations

Equation 2 performs multiple regression, regressing each *X_j_*(*t* + Δ*t*) onto each *X_j_*(*t*) whilst controlling for all other state estimates at time *t*. This method works well when spurious relationships between *X_i_*(*t*) and *X_j_*(*t* + Δ*t*) are mediated by the subspace spanned by the other estimated states at time t (in particular *X_j_*(*t*)). One situation in which this assumption might be challenged is when replay is superimposed on a large neural oscillation. For example, during rest with eyes closed (which is often of interest in replay analysis), MEG and EEG data often express a large alpha rhythm, at around 10Hz.

If all states experience the same oscillation at the same phase, the approach correctly controls false positives. The oscillation induces a spurious correlation between *X_i_*(*t*) and *X_j_*(*t* + Δ*t*) but, as before, this spurious correlation is mediated by *X_j_*(*t*).

However, this logic fails when states experience the oscillation at different phases. This scenario may occur, for example, if there are travelling waves in cortex ^19,20^, because different sensors will experience the wave at different times, and different states have different contributions from each sensor. In this case, *X_i_*(*t*) predicts *X_j_*(*t* + Δ*t*) over and above *X_j_*(*t*).

To see this, consider the situation where Δ*t* is 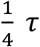 (where *τ* is the oscillatory period) and the phase shift between *X_i_*(*t*) and *X_j_*(*t*) is pi/2. Now every peak in *X_j_*(*t* + Δ*t*) corresponds to a peak in *X_i_*(*t*) but a zero of *X_j_*(*t*).

To combat this problem, we can include phase shifted versions/more timepoints of *X(t*). If dominant background oscillation is at alpha frequency (e.g., 10Hz), neural activity at time T would be correlated with activity at time T + *τ*. We can control for that, by including *X*(*t + τ*), as well as *X*(*t*) in the GLM (Fig. 3b). Here *τ* = 100 ms, if assuming the frequency is 10Hz. Applying this method to the real MEG data during rest, we see much diminished 10Hz oscillation in sequence detection during rest ^12^.

#### Spatial correlations

As mentioned above, correlations between decoded variables commonly occur. The simplest type of decoding model is a binary classifier that maps brain activity to one of two states. These states will, by definition, be perfectly anti-correlated. Conversely, if separate classifiers are trained to distinguish each state’s representation from baseline (“null”) brain data, then the states will often be positively correlated with each other.

Unfortunately, positive or negative correlation between states reduces the sensitivity of sequence detection, because it is difficult to distinguish between states within the sequence: collinearity impairs estimation of *β* in Equation 2. In Figure 3c, we show in simulation that the ability to detect real sequences goes down as spatial correlation goes up.

Ideally, the state decoding models should be as independent as possible. We have suggested the approach of training models to discriminate one state against a mixture of other states and null data ^11,12^. The mixture ratio can be adjusted. Adding more null data causes the states to be positively correlated with each other, while less null data leads to negative correlation. We adjust the ratio to bring the correlation between states as close to zero as possible. In Figure 3d, we show in simulation the benefit for sequence detection. An alternative method is penalizing covariance between states in the classifier’s cost function ^21^.

#### Regularization

A key parameter in training high dimensional decoding models is the degree of regularization.

In sequence analysis, we are often interested in spontaneous reactivations of state representations - as in replay. However, our decoding models are typically trained on stimulus-evoked data, because this is the only time at which we know the ground truth of what is being represented. This poses a challenge in so far as the models best suited for decoding evoked activity at training may not be well suited for decoding spontaneous activity at subsequent test.

We find that L1 weight regularization outperforms L2 regularization in detecting sequences (Fig. 3e). Notably, the L1 penalty encourages sparsity, which reduces spatial correlation between states.

### STATISTICAL INFERENCE

So far, we have shown how to quantify sequences in representational dynamics. An essential final step is assessing the statistical reliability of these quantities.

All the tests described in this section evaluate the consistency of sequences *across subjects*. This is very important, because even in the absence of any real sequences of task-related representations, spontaneous neural activity is not random but follows repeating dynamical motifs ^22^. Solving this problem requires a randomized mapping between the assignment of physical stimuli to task states. This can be done across subjects, permitting valid inference at the group level.

At the group level, the statistical testing problem can be complicated by the fact that sequence measures do not in general follow a known distribution. Additionally, if the state-to-state lag of interest (Δ*t*) is not known a priori, it will be necessary to perform tests at multiple lags, creating a multiple comparisons problem over a set of tests with complex interdependencies. In this section we discuss inference with these issues in mind.

#### Distribution of sequenceness at a single lag

If the state-to-state lag of interest (Δ*t*) is known a priori then the simplest approach is to compare the sequenceness against zero, for example using either a signed-rank test, or one-sample *t* test (assuming Gaussian distribution). Such testing assumes that the data would be centred on zero if there were no real sequences. We show this approach is safe in both simulation (assuming no real sequences) and real MEG data in which we know there are no sequences.

In simulation, we assume no real sequences, but state time courses are autocorrelated. At this point, there is no systematic structure in the correlation between the neuronal representations of different states (see later for this consideration). We then simply select the 40 ms time lag and compare its sequenceness to zero using either a signed-rank test or one-sample *t* test. We compare false positive rates predicted by the statistical tests with false positive rates measured in simulation (Fig. 4a). We see the empirical false positives are well predicted by theory.

**Fig. 4.**
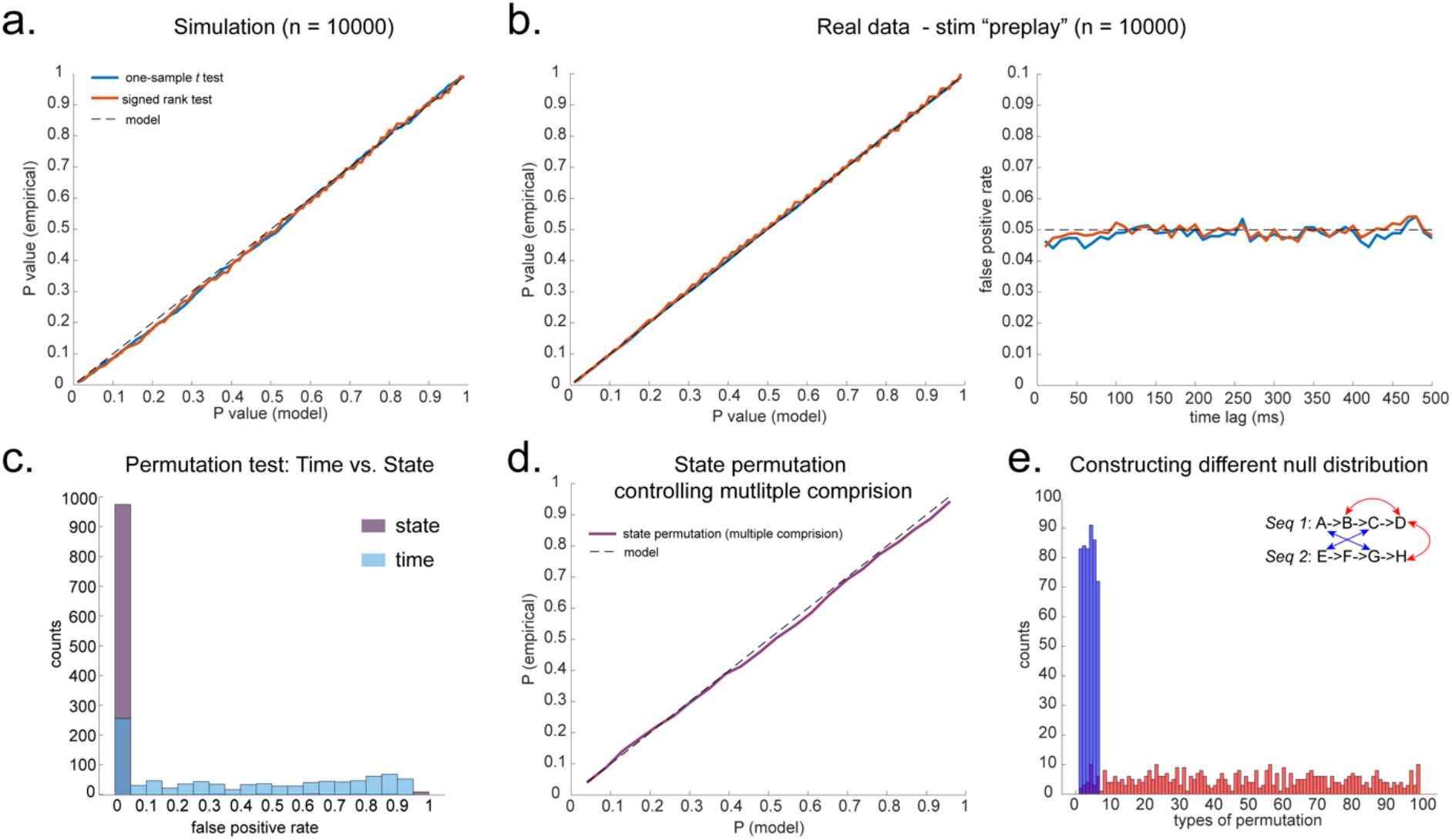
Statistical inference. **a,** P-P plot of one-sample *t* test (blue) and Wilcoxon signed rank test (red) against zero. This is done in simulated MEG data assuming auto-correlated state time courses but no real sequences. In each simulation, the statistics are done only on sequenceness at 40 ms time lag, across 24 simulated subjects. There are 10,000 simulations. **b,** We have also tested the sequenceness distribution on the real MEG data. This is pre-task resting state of 22 subjects from Liu et. al, where the ground truth is no sequence given the stimuli are not shown yet. The statistics are done on sequenceness at 40 ms time lag, across the 22 subjects. There are eight states. The state identity is randomly shuffled 10,000 times to construct the null distribution. **c,** Time-based permutation test tends to give high false positive, while state identity-based permutation does not. This is done in simulation assuming no real sequences (n=1000). **d,** P-P plot of state identity-based permutation test over peak sequenceness is shown. To control for multiple comparisons, the null distribution is formed taking the maximal absolute value over all computed time lags within a permutation, and the permutation threshold is defined as the 95% percentile over permutations. In simulation, we only compared the max sequence strength in the data to this permutation threshold. There are 10,000 simulations. In each simulation, there are 24 simulated subjects, with no real sequence. **e,** In state-identity based permutation, we can test more specific hypotheses by controlling the null distribution. Blue are the permutations that only exchange state identity across sequences. Red are the permutations that permit all possible state identity permutations. 500 random state permutations are chosen from all possible ones. The X axis is the different combinations of the state permutation. It is sorted so that the cross-sequence permutations are in the beginning.

**Fig. 5.**
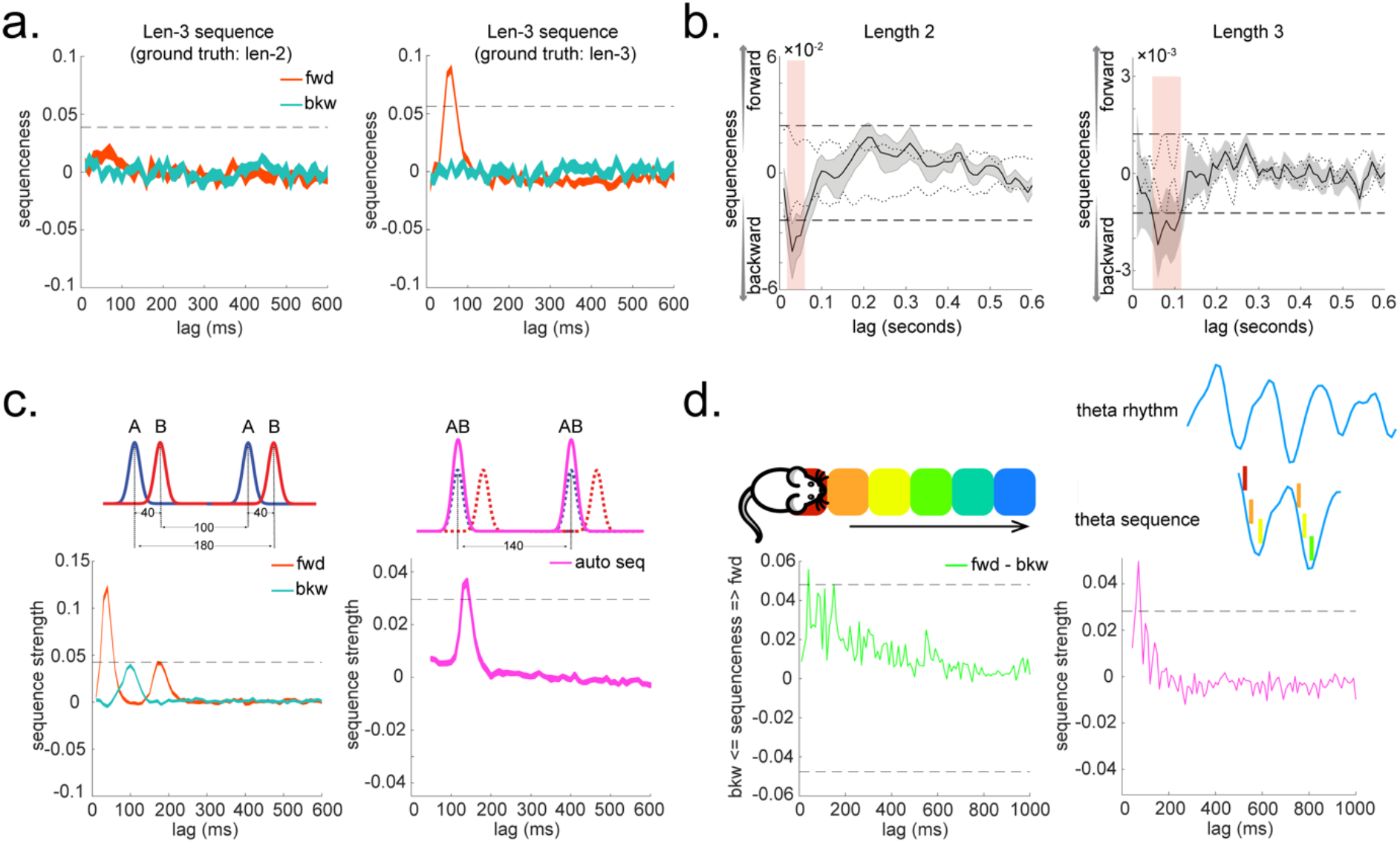
Extension to TDLM: Multi-step sequences and Sequence of sequences. **a**, TDLM can quantify not only pair-wise transition, but also longer length sequences. It does so by controlling for evidence of shorter length to avoid false positive. **b,** Method applied to human MEG data, incorporating control of both alpha oscillation and co-activation for both length-2 and length-3 sequence length. Dashed line indicates the permutation threshold. This is adapted from Liu, et al. ^11^. **c,** TDLM can also be used iteratively to capture the repeating pattern of sequence event itself. Illustration in the top panel describes the ground truth in the simulation. Intra-sequence temporal structure (right) and intersequence temporal structure (right) can be extracted simultaneously. **d,** On a real rodent hippocampal electrophysiological dataset, TDLM revealed the well-known theta sequence phenomena during active spatial navigation.

We also tested this on real MEG data. In Liu, et al. ^12^ we had one condition where we measured resting activity before the subjects saw any stimuli. Therefore, by definition these stimuli could not replay, but we can use the classifiers from these stimuli (measured later) to test the false positive performance of statistical tests on replay. To get many examples, we randomly permute the 8 different stimuli 10,000 times and then compare sequenceness (at 40 ms time lag) to zero using either signed rank test or one-sample *t* test across subjects. Again, predicted and measured false positive rates match well (Fig. 4b, left panel). This holds true across all computed time lags (Fig. 4b, right panel).

An alternative to making assumptions about the form of the null distribution is to compute an empirical null distribution by permutation. Given that we are interested in the sequence of states over time, one could imagine permuting either state identity or time. However, permuting time uniformly will typically lead to a very high incidence of false positives, as time is not exchangeable under the null hypothesis (Fig. 4c, blue colour). Permuting time destroys the temporal smoothness of neural data, creating an artificially narrow null distribution ^11,12^. State permutation, on the other hand, only assumes state identities are exchangeable under the null hypothesis, while preserving the temporal dynamics of the neural data, represents a safer statistical test that is well within 5% false positive rate (Fig. 4c, purple colour).

#### Correcting for multiple comparisons

If the state-to-state lag of interest is not known, we have to search over a range of time lags. As a result, we have a multiple comparison problem. Unfortunately, we don’t yet have a good parametric method to control for multiple testing over a distribution. It is possible that one could use methods that exploit the properties of Gaussian Random Fields, as is common in fMRI ^23^, but we have not evaluated this approach. We could use Bonferroni correction, but the assumption that each computed time lag is independent is likely false and overly conservative.

We recommend relying on state-identity based permutation. To control the family wise error rate (assuming *α* = 0.05), we want to make sure that there is a 5% probability of getting the tested sequenceness strength (*S_test_*) or bigger by chance in *any* of the multiple tests. We therefore need to know what fraction of the permutations give *S_test_* or bigger in any of *their* multiple tests. If any of the sequenceness scores in each permutation exceed *S_test_*, then the ***maximum*** sequenceness score in the permutation will exceed *S_test_*, so it is sufficient to test against the maximum sequenceness score in the permutation. The null distribution is therefore formed by first taking the peak of sequenceness across all computed time lags of each permutation. This is the same as approach as is used for family-wise error correction for permutations tests in fMRI data ^24^, and in our case it is shown to behave well statistically (Fig. 4d).

#### What to permute

We can choose which permutations to include in the null distribution. For example, consider a task with two sequences, *Seq1: A → B → C → D*, and *Seq2: E → F → G → H*. We can form the null distribution either by permuting all states (e.g., one permutation might be: E→ *F → A → B*, H→ *C → E → D*), as was performed in Kurth-Nelson, et al. ^11^. Alternatively, we can form a null distribution which only includes transitions between states in different sequences (e.g., one permutation might be: D→ *G → A → E*, H→ *C → F → B*), as was performed in Liu, et al. ^12^. In each case, permutations are equivalent to the test data under the assumption that states are exchangeable between positions and sequences. The first case has the advantage of many more possible permutations, and therefore may make more precise inferential statements in the tail (Fig 4e). The second may be more sensitive in the presence of signal, as the null distribution is guaranteed not to include permutations which share any transitions with the test data.

#### Cautionary note on exchangeability of states after training

Until now, all tests have assumed that state identity is exchangeable under the null hypothesis. Under this assumption, it is safe to perform state-identity based permutation tests on *Z_F_* and *Z_B_*. In this section, we consider a situation where this assumption is broken.

More specifically, we are considering a situation where the neural representation of state *A* and *B* are related in a systematic way or, in other words, the classifier on state *A* is confused with state *B*, and we are testing sequenceness of *A → B*. Crucially, to break the exchangeability assumption, representations of *A* and *B* have to be systematically more related than other states, e.g., *A* and *D*. This cannot be caused by low level factors (e.g., visual similarity) because states are counterbalanced across subjects, so any such bias would cancel at the population level. However, such a bias might be *induced* by task training.

In this situation, it is, in principle, possible to detect sequenceness of *A → B*, even in the absence of real sequences. In the autocorrelation section above, we introduced protections against the interaction of state correlation with autocorrelation. These protections may fail in the current case as we cannot use other states as controls (as we do in the multiple linear regression), because *A* has systematic relationship with *B*, but not other states. State permutation will not protect us from this problem because state identity is no longer exchangeable.

Is this a substantive problem? After extensive training, behavioural pairing of stimuli can indeed result in increased neuronal similarity ^25,26^. These early papers involved long training in monkeys. More recent studies have shown induced representational overlap in human imaging within a single day ^27–29^ However, when analysed across the whole brain, such representational changes tend to be localised to discrete brain regions ^30,31^, and as a consequence may have limited impact on whole brain decodeability.

Whilst we have not yet found a simulation regime in which false positives are found (as opposed to false negatives), there exists a danger in cases where, by experimental design, the states are not exchangeable.

### EXTENSIONS TO TDLM

TDLM can be used iteratively. Two extensions of TDLM of particular interest are: Multi-step sequences and Sequence of sequences. The former asks about consistent regularity among multiple states, the latter ask about the hierarchical structure of state reactivation, not only within but between sequences.

#### Multi-step sequences

So far, we have introduced methods for quantifying the extent to which the state-to-state transition structure in neural data matches a hypothesized task-related transition matrix. An important limitation of these methods is that they are blind to hysteresis in transitions. In other words, they cannot tell us about multi-step sequences. In this section, we describe a methodological extension to measure evidence for sequences comprising more than one transition: for example, *A → B → C*.

The key ingredient is controlling for shorter sub-sequences (e.g., *A → B* and *B → C*), in order to find evidence unique to the multi-step sequence of interest.

Assuming constant state-to-state time lag, Δ*t*, between A and B, and between B and C. We can create new state space AB, by shifting B up Δ*t*, and elementwise multiply it with state A. This new state AB measure the reactivation strength of *A → B*, with time lag Δ*t*. In the same way, we can create new state space, BC, AC, etc. Then we can construct the same first level GLM on the new state space. For example, if we want to know the evidence of *A → B → C* at time lag Δ*t*. We can regress AB onto state time course C, at each Δ*t* (cf. Equation 1). But we want to know the unique contribution of AB to C. More specifically, we want to test if the evidence of *A → B → C* is stronger than *X → B → C*, where X is any state but not A. Therefore, similar as Equation 2, we want to control CB, DB, when looking for evidence of AB of C. Applying this method, we show TDLM successfully avoids false positives arising out of strong evidence for shorter length (see simulation results in Fig. 3f, see results obtained on human neuroimaging data in Fig. 3g). This process can be generalized to any number of steps.

#### Sequence of sequences

We have so far detailed use of either sensory or abstract representations as states in TDLM. We now take a step further and use sequences themselves as states. With this kind of hierarchical analysis, we can search for sequences of sequences. This is useful because it can reveal the temporal structure not only within sequence, but also between sequences. The organization between sequences is of particular interest for revealing neural computations. For example, the forward and backward search algorithms hypothesized in planning and inference ^32^ can be cast as sequences of sequences problem: the temporal structure of forward and backward sequence. This can be tested by using TDLM iteratively.

As yet little human neural data is available on the organization of sequences. Interestingly, one can think of theta sequence, a well-documented phenomenon during rodent spatial navigation ^8,13,33^, as a neural sequence repeating itself in theta frequency (6 - 12 Hz). We will show TDLM is able to replicate this well-known phenomenon.

To look for sequences between sequences we need first to define sequences as new states. To do so, the raw state course, for example, state B needs to be shifted up by the empirical within-sequence time lag Δ*t* (determined by the two-level GLM described above), to align with the onset of state A, if assuming sequence *A → B* exist (at time lag Δ*t*). Then, we can elementwise multiply the raw state time course A with the shifted time course B, resulting in a new state AB (Fig. 2d). Each entry in this new state time course indicates the reactivation strength of sequence AB at a given time.

After that, the general two-level GLMs framework still applies, but with one important caveat. The new sequence state (e.g., AB) is defined based on the original states (A and B), and we are now interested in the reactivation regularity, i.e., sequence, between sequences, rather than the original states. We should therefore control for the effects of the original states. Effectively, this is like controlling for main effects (e.g., state A and shifted state B) when looking for their interaction (sequence AB). TDLM achieves this by putting time lagged original state regressors A, B, in addition to AB, in the first level GLM sequence analysis (see details in online Methods).

In simulation we demonstrate, applying this method, that TDLM can uncover hierarchical temporal structure: state A is temporally leading state B with 40 ms lag, and the sequence A->B tends to repeat itself with a 140 ms gap (Fig. 4a). On real rodent hippocampal electrophysiological recording, we replicate the well-known theta sequence - neural sequence repeating itself in theta frequency (Fig. 4b, see detailed analysis on rodent data in Supplementary Note 2).

In addition to looking for temporal structure of the same sequence, this method is equally suitable when searching for temporal relationship between difference sequences in a general form. For example, assuming two different types of sequences, one sequence type has a within-sequence time lag at 40 ms; while the other has a within-sequence time lag at 150 ms; and there is a gap of 200 ms between the two types of sequences (Supplementary Fig. 3a) (these time lags are set arbitrarily for illustration purposes. TDLM captures accurately the dynamics both within and between the sequences (Supplementary Fig. 3b, c), supporting a potential for uncovering temporal relationships between sequences in general under the same framework.

### SOURCE LOCALIZATION

Uncovering the temporal structure of neural representation is important, but one might also want to ask where in the brain the sequence is generated. Rodent electrophysiology research focuses mainly on hippocampus when searching for replay. One advantage of whole-brain non-invasive neuroimaging over electrophysiology (despite many known disadvantages, including poor anatomical precision, low signal-noise ratio) is its ability to look for neural activity in other brain regions. Ideally, we would like a method that is capable of localizing sequences of more abstract representation in brain regions beyond hippocampus ^12^.

We can achieve this by availing of the two-level GLMs in TDLM. More specifically, after identifying the empirical time lag that gives rise to the strongest neural sequence, one can project the time lag back to the time series of decoded states and work out the probability of sequence reactivation at each time point (Supplementary Fig. 4a, left panel). This is the same as changing the state space from A and B to sequence A->B. This gives us a temporal stamp on the testing time, e.g., resting state: the time indices of sequence onset (Supplementary Fig. 4a, right panel). To ensure it is the onset of a sequence event, rather than a middle portion, we apply an extra constraint: there is a low sequence probability time window (e.g., 200 ms) before the sequence onset. Then, we can epoch the testing time data into a bunch of sequence events with appropriate thresholding, e.g., above 95^th^ percentile (see detailed calculation in online Method). After epoching, the epoched data can be treated as event related neural activity, with onset as the initialization of neural sequence. This approach is similar to spike-triggered averaging ^34,35^. Applying this to real MEG data during rest, we can detect increased hippocampal power at 120-150 Hz, during replay onset (Supplementary Fig. 4b, c).

## DISCUSSION

TDLM is a general analysis framework for capturing sequence regularity of neural representations. It is developed on human neuroimaging data but can be applied to other data sources, including rodent electrophysiology recordings. The framework can facilitate crossspecies investigations and enables investigation of phenomena that are not readily addressable in rodents ^12^.

The temporal dynamics of neural states have been studied previously with MEG ^22,36^. Normally states are defined by common physiological features (e.g., frequency, functional connectivity) during rest, and termed resting state networks (e.g., default mode network ^37^). However, these approaches remain agnostic about the ***content*** of neural representation. Being able to study the temporal dynamics of *representational content* permits richer investigations into cognitive processes, as neural states can be analysed in the context of their roles with respect to cognitive tasks.

Reactivation of neural representations have also been studied previously ^38^ using approaches similar to the decoding step of TDLM, or multivariate pattern analysis (MVPA) ^39^. This has proven fruitful in revealing mnemonic functions^29^, understanding sleep^40^, and decisionmaking^41^. However, classification alone cannot reveal the rich temporal structures of reactivation dynamics. For example, the ability to detect sequences allows us to tease apart clustered from sequential reactivation, where this may be important for dissociating decision strategies ^42^ and their individual differences ^42,43^. Furthermore, it enables comparisons with the sequential reactivation patterns reported in rodent hippocampus ^10,44^, and may allow tests of neural predictions from process models such as reinforcement learning ^45^, which have been hard to probe previously in humans ^46^.

We have mainly discussed the application of TDLM on high temporal resolution neuroimaging data (e.g., MEG). Recently, sequential replay has been reported using fMRI ^47^. We anticipate it will be useful to combine the high temporal resolution available in M/EEG and the spatial precision available in fMRI to probe region - specific sequential computation. Whilst related techniques are available ^48^, TDLM could, in principle, also be applied to fMRI data.

TDLM enables neuroscientists to decipher rich temporal structures of neural reactivation. We described the application of TDLM mostly during off-task state. However, the very same analysis can be applied to on-task data, to test for cued sequential reactivation ^43^, or sequential decision-making ^46^. We believe TDLM opens doors for novel investigations of human cognition, including language, sequential planning and inference in non-spatial cognitive tasks ^11,42^. It is particularly suited to test specific neural prediction from process models. Therefore, we hope TDLM can aid a synthesis between empirical and theoretical approaches in neuroscience and in so doing shed novel lights on dynamic neural computation.

## ONLINE METHODS

### The TDLM framework

TDLM comprises three stages. The objective of the first stage is to map the multivariate neural activity to labelled states. In the second and third stage, TDLM builds a linear generative model of the state time course and assesses the statistical significance by non-parametric permutation test. The three-stage computation is described below.

#### Step 1: Mapping multivariate neural activity to labelled state

Mappings between neural activity and labelled states are established through a supervised decoding approach. To avoid selection bias, the training data should come from an independent task with no biased experience of the states. The choice of machine learning methods is not the focus of this work. We choose logistic regression models for simplicity.

For each state we trained one binomial classifier. Positive examples for the classifier were trials on which that object was presented. Negative examples consisted of two kinds of data: trials when another object was presented, and data from the fixation period before the semantic precue appeared (i.e., “null data”). The null data are included to reduce the anticorrelation between different classifiers. It was possible for all classifiers to report low probabilities simultaneously in testing data. Prediction accuracy was estimated by treating the highest probability output among all classifiers as the predicted object. Permutation-based method is employed to assess the statistical significance, which is corrected for multiple comparisons across time. The time point that gives the highest cross-validation accuracy is selected for training the classifiers.

#### Step 2: Quantifying strength of state transitions

The decoding models allow one to measure spontaneous reactivation of task-related representations during testing time, where ground truth label is not available e.g., resting state. TDLM defined a ‘sequenceness’ measure, which describes the degree to which these representations were reactivated in a prescribed sequential order.

TDLM first applies each of the state decoding models to the test data. This yields a reactivation matrix X with dimension *T × N*., where *T* is the duration of the time series, and *N* is the number of states. After that, TDLM asks whether particular sequences of state activations appeared above chance in the reactivation matrix X, by applying a two-level GLMs (Equations 2 and 5).

#### Step 3: Test statistical significance of neural sequence

In the final step, TDLM assesses the statistical significance of sequenceness. TDLM chooses to apply nonparametric tests involving possible permutations of the state labels. Shuffling state identity allows one to reject the null hypothesis that the MEG time series had no relationship to the transition structure of the task. Different levels of statistical inference can be made precisely by controlling the null distribution - how the states are permuted.

### Change of state spaces

TDLM allows one to construct a new state space by building on the original one. For example, after figuring out the empirical state-to-state time lag, TDLM can build the sequence state, and look for sequence of sequence, or identify the onset of sequence.

To do so, TDLM first time shifted the reactivation matrix X up to the empirical time lag *Δt_b_*, obtained *X_Δb_*.

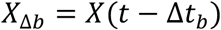

Then, *X* is multiplied by the transition matrix *P*, obtained a project matrix - *X_P_*.

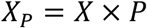

Next, TDLM elementwise multiply *X_Δb_* by *X_P_*, result in matrix *R*, which indicate the new state space of sequence, where each element indicates the strength of a (pairwise) sequence at a given moment in time.

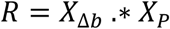

### Identifying sequence onsets

Sequence onsets were defined as moments when a strong reactivation of a state was followed by a strong reactivation of the next state in the sequence. In general, TDLM first finds the empirical state-to-state time lag *Δt_b_* where there is maximum evidence for state-to-state sequenceness. Finally, TDLM identifies the sequence onset by thresholding the sequence state at its high (e.g., 95^th^) percentile with a constraint that a sequence onset has a sequence-free time window (e.g., 100 ms) preceding it. This analysis pipeline gives a temporal stamp on the testing time. One can therefore epoch the data based on those sequence onsets and apply temporal frequency analysis and source localization, just like on the standard task data.

### Sequence of sequence

TDLM is capable of quantifying not only the item-to-item transitions, but also sequence-to-sequence dynamics after change of state space. To quantify sequence of sequences, TDLM needs to construct the design matrix to carefully control for dynamics within the sequence. In the linear model, this is effectively asking for the interaction effect of item state A and B, one should therefore control for the main effect of A and B. Similar with quantifying the original state-to-state transitions, TDLM operate in two-level GLMs to measure the sequence-to-sequence transitions, but with extra control of within sequence effects.

Let’s assume the sequence state matrix is *X_seq_*, after transforming the original state space to sequence space based on the empirical within-sequence time lag *Δt_w_*. Each column at *X_seq_* is sequence state, denoted by *S_ij_*, which indicates the strength of sequence *i* -> *j* reactivation. The raw state *i* is *X_i_*, and the shifted raw state *j* is *X_jw_* (by time lag *Δt_w_*).

In the first level GLM, TDLM ask for the strength of unique contribution of sequence state *S¿j* to *S_mn_* while controlling for original states (*X¿* and *X_JW_*). For each sequence state *ij*, at each possible time lag Δ*t*, TDLM estimated a separate linear model:

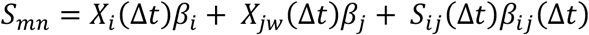

Repeat this process for each sequence state separately at each time lag, resulting a sequence matrix *β_seq_*.

In the 2^nd^ level GLM, TDLM asks how strong the evidence of sequence of interest is compared to sequences that have the same starting state or end state at each time lag. *β_seq_* contains the beta of sequence of interest, *β_sij_*; beta of sequence that share the same starting state, *β_si_*; beta of sequence that share the same end state, *β_Sj_*. The design matrix *T* includes sequence of sequence of interest *T_seq_* (label “1” for *β_ij__Δst_*, “0” for anything else), constant, *T_C_*.

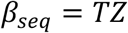

This is asking how strong the specific sequence of sequence of interest (*T_seq_*) is, compared to the general sequence effect that is contributed by either the same starting state or the end state. This resulted in a single number *Z_seq_*, quantifying the specific strength of transition of interest at given time lag Δ*t*.

### Abstract code

TDLM assumes the variance of structure code and sensory code of the same object are uncorrelated and can be linearly decomposed. TDLM first estimates the mean multivariate response pattern of the objects (sensory code) on the data where there is no structural information. The objective here is to find the response patterns that can explain the maximal variance of sensory code overall rather than separating neural representations of each sensory code ^49^. After that, TDLM regresses the multivariate response pattern of sensory code, *β_sensory_*, onto the position code training data, *D_p_*, and get the residual that cannot be explained by the sensory code, *E_P_*.

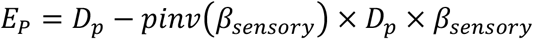

The position code is then trained only on the residuals, *E_P_*, through the same analysis pipeline of the first step of TDLM.

### Human MEG dataset

#### Task design

Participants were required to perform a series of tasks with concurrent MEG scanning (see details in Liu, et al. ^12^). The functional localizer task was performed before the main task and was used to train a sensory code for eight distinct objects. Note, the participants were provided with no structural information at the time of the localizer. These decoding models, trained on the functional localizer task, capture a sensory level neural representation of stimuli (i.e.,stimulus code). Following that, participants were presented with the stimuli and were required to unscramble the “visual sequence” into a correct order, i.e., the “unscrambled sequence” based on a structural template they had learned the day before. After that, participants were given a rest for 5 mins. In the end, stimuli were presented again in random order, and participants were asked to identify the true sequence identity and structural position of the stimuli. Data in this session are used to train the structural code of the objects.

#### MEG data Acquisition and Pre-processing

MEG was recorded continuously at 600 samples/second using a whole-head 275-channel axial gradiometer system (CTF Omega, VSM MedTech), while participants sat upright inside the scanner. Participants made responses on a button box using four fingers as they found most comfortable. The data were resampled from 600 to 100 Hz to conserve processing time and improve signal to noise ratio. All data were then high-pass filtered at 0.5 Hz using a first-order IIR filter to remove slow drift. After that, the raw MEG data were visually inspected, and excessively noisy segments and sensors were removed before independent component analysis (ICA). An ICA (FastICA, http://research.ics.aalto.fi/ica/fastica) was used to decompose the sensor data for each session into 150 temporally independent components and associated sensor topographies. Artefact components were classified by combined inspection of the spatial topography, time course, kurtosis of the time course and frequency spectrum for all components. Eye-blink artefacts exhibited high kurtosis (>20), a repeated pattern in the time course and consistent spatial topographies. Mains interference had extremely low kurtosis and a frequency spectrum dominated by 50 Hz line noise. Artefacts were then rejected by subtracting them out of the data. All subsequent analyses were performed directly on the filtered, cleaned MEG signal, in units of femtotesla.

#### MEG Source Reconstruction

All source reconstruction was performed in SPM12 and FieldTrip. Forward models were generated on the basis of a single shell using superposition of basis functions that approximately corresponded to the plane tangential to the MEG sensor array. Linearly constrained minimum variance beamforming ^50^, was used to reconstruct the epoched MEG data to a grid in MNI space, sampled with a grid step of 5 mm. The sensor covariance matrix for beamforming was estimated using data in either broadband power across all frequencies or restricted to ripple frequency (120-150 Hz). The baseline activity was the mean neural activity averaged over −100 ms to −50 ms relative to sequence onset. All non-artefactual trials were baseline corrected at source level. We looked at the main effect of the initialization of sequence. Non-parametric permutation tests were performed on the volume of interest to compute the multiple comparison (whole-brain corrected) P-values of clusters above 10 voxels, with the null distribution for this cluster size being computed using permutations (n = 5000 permutations).

## Code availability

Source code of TDLM can be found at https://github.com/yunzheliu/TDLM.

## Data availability

Data are available upon reasonable request from the corresponding author, unless prohibited owing to research participant privacy concerns.

## Acknowledgement

We thank Matthew A. Wilson for providing rodent data and help with the rodent data analysis. We thank Matt Nour and Mark Woolrich for helpful comments on a previous version of the manuscript. Y.L. is also grateful for the unique opportunity provided by the Brains, Minds & Machines Summer Course. We acknowledge funding from UCL Graduate Research Scholarship and Overseas Research Scholarship to Y.L., Wellcome Trust Investigator Award (098362/Z/12/Z) to R.J.D., and Wellcome Trust Senior Research Fellowship (104765/Z/14/Z), and Principal Research Fellowship (219525/Z/19/Z), together with a James S. McDonnell Foundation Award (JSMF220020372), to T.E.J.B. Both Wellcome Centres are supported by core funding from the Wellcome Trust: Wellcome Centre for Integrative Neuroimaging (203139/Z/16/Z), Wellcome Centre for Human Neuroimaging (091593/Z/10/Z). The Max Planck UCL Centre is a joint initiative supported by UCL and the Max Planck Society.

## SUPPLEMENTARY FIGURES

**Supplementary Fig. 1.**
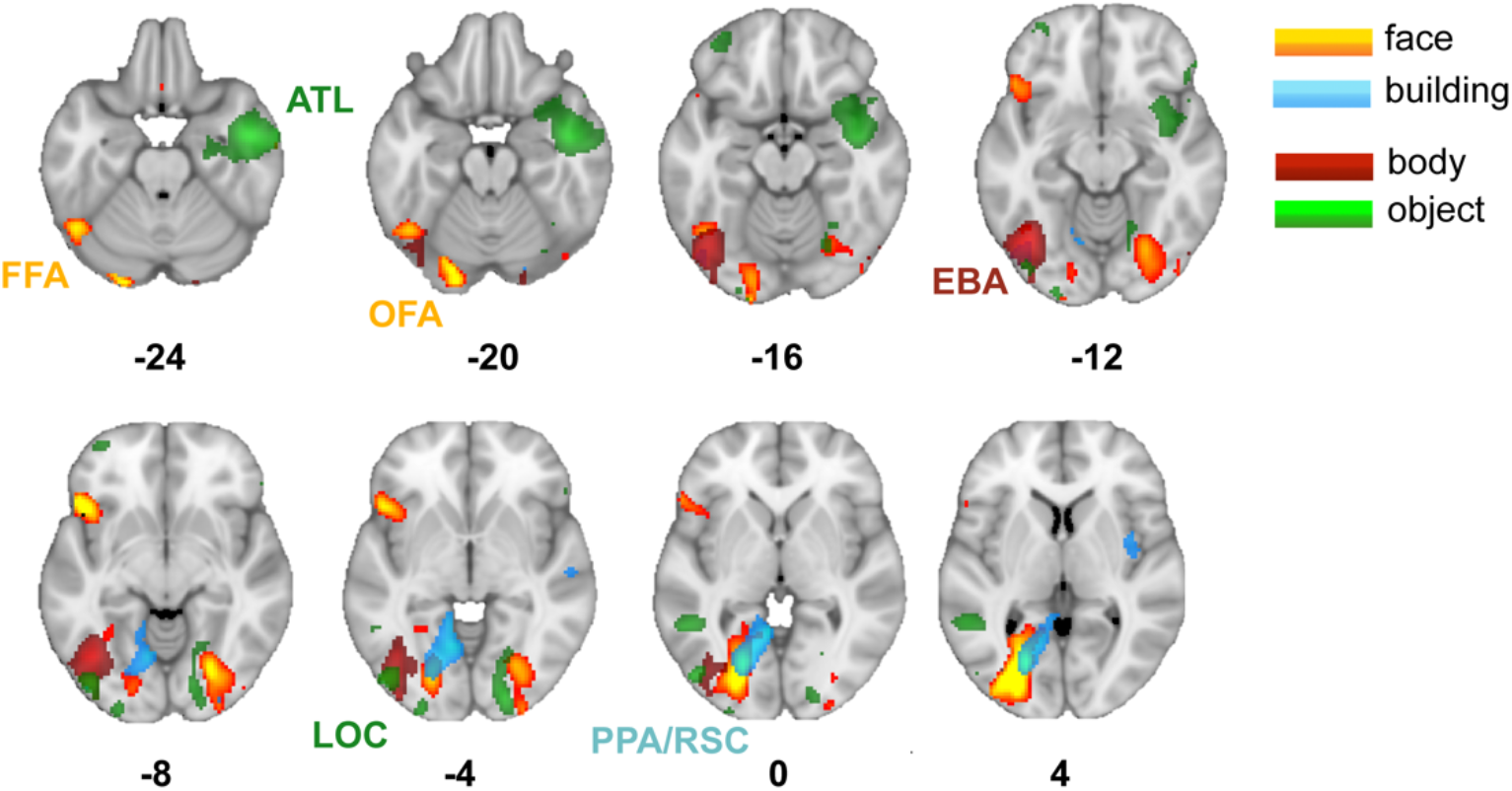
Source localization of stimuli evoked neural activity. The states here are defined as the stimuli evoked neural activity. The classifiers are trained at 200 ms post-stimulus onset. For example, the stimuli are faces, buildings, body parts, and objects. Source localizing the evoked neural activity, we found expected activation pattern of the 4 stimuli based on literature. For faces, activation peaked in a region roughly consistent with the fusiform face area (FFA) as well as the occipital face area (OFA). Activation for building stimuli was located between the well-known parahippocampal place area (PPA) and the retrosplenial cortex (RSC), a region also known to respond to scene and building stimuli. Activation for body part stimuli was in a region consistent with the extrastriate body area (EBA). Activation for objects was in a region consistent with the object-associated lateral occipital cortex (LOC) as well as an anterior temporal lobe (ATL) cluster that may relate to conceptual processing of objects. Individual category maps thresholded to display localized peaks for illustration. This is adapted from Wimmer, et al. ^2^. Full unthresholded maps can be found at https://neurovault.org/collections/6088/.

**Supplementary Fig. 2.**
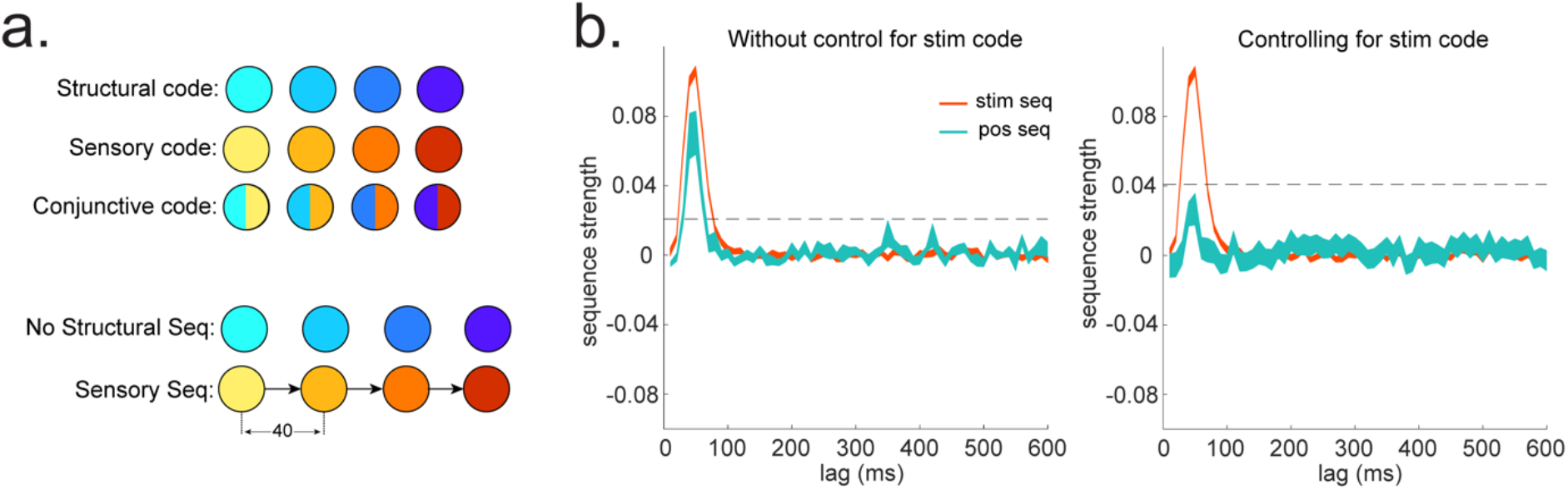
Sequences of abstract code. **a,** Illustration of the relationship between sensory code and (abstract) structural code. The problem is we cannot directly access structural code. We can only indirectly obtain structural code from the conjunctive code which have both sensory and structural information. In the ground truth, there is sequence of sensory code but not structural code. **b,** We show in simulation the importance of controlling for sensory (stim) information, when looking for sequences of abstract code.

**Supplementary Fig. 3.**
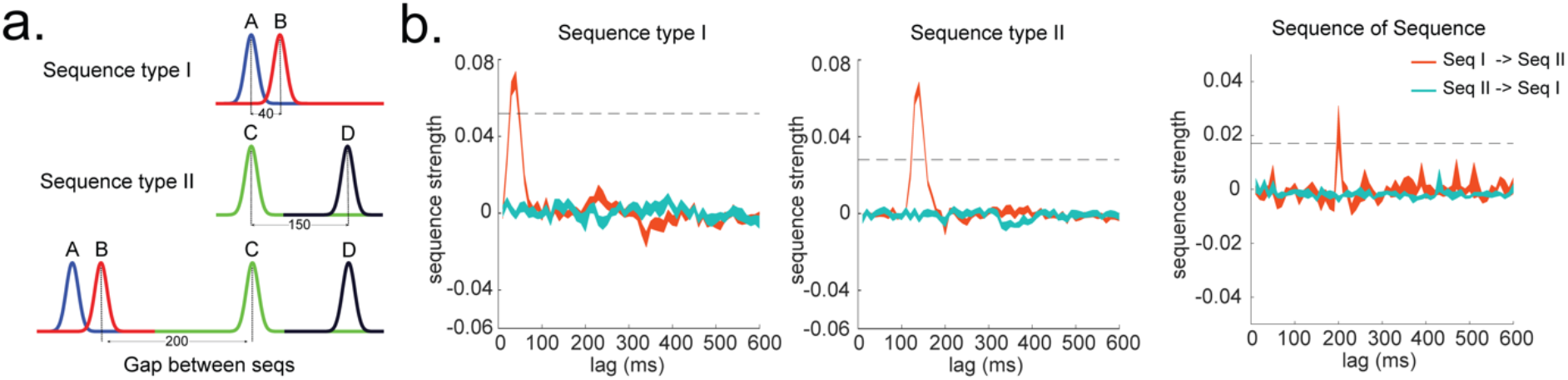
Temporal structure between and within different sequences. **a,** Illustration of two sequence types with different state-to-state time lag within sequence, and a systematic gap between the two types of sequences. **b,** TDLM can capture the temporal structures both within (left panel) and between (right panel) the two sequence types.

**Supplementary Fig. 4.**
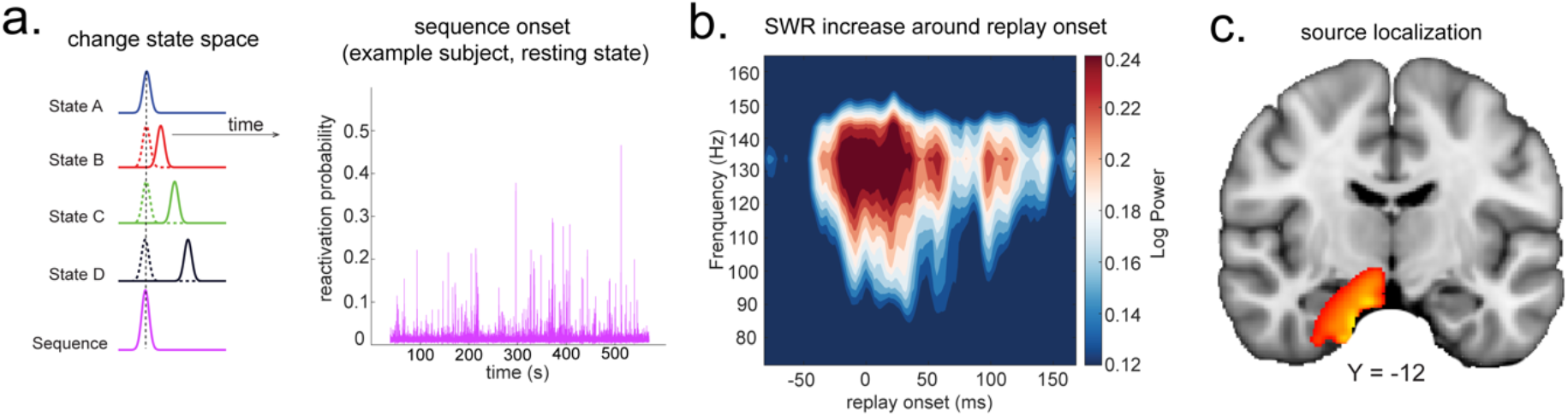
Source localization of replay onset. **a,** TDLM figures out the onset of sequence based on the identified optimal state-to-state time lag (left panel). Sequence onset during resting state from one example subject is shown (right panel). **b,** On the real human neuroimaging data during rest, there was a significant power increase (averaged across all sensors), in the ripple frequency band (120-150 Hz), at the onset of replay, compared to the pre-replay baseline (100 to 50 ms before replay). **c,** Source localization of ripple-band power at replay onset revealed significant hippocampal activation (right panel, peak MNI coordinate: X = 18, Y = −12, Z = −27). This is adapted from Liu, et al. ^1^.

## Supplementary Note 1: Synthetic datasets

We simulate the data to be similar with the human MEG data. Therefore, there is always strong autocorrelation in time, and sometimes with rhythmic oscillation (e.g., 10Hz). In spatial domain, the sensors are always spatially correlated. To imitate the temporal auto-correlation feature of real MEG data, the simulated data is generated with an auto-aggressive model with multivariate gaussian noise. To imitate the spatial correlation between MEG sensors, we add dependence among sensors in the simulated data. The simulated data is nSensors (number of sensors = 273) by nSamples (number of samples = 6000, with each sample =10 ms).

An example of the Matlab implementation are (see full codebase in https://github.com/yunzheliu/TDLM):

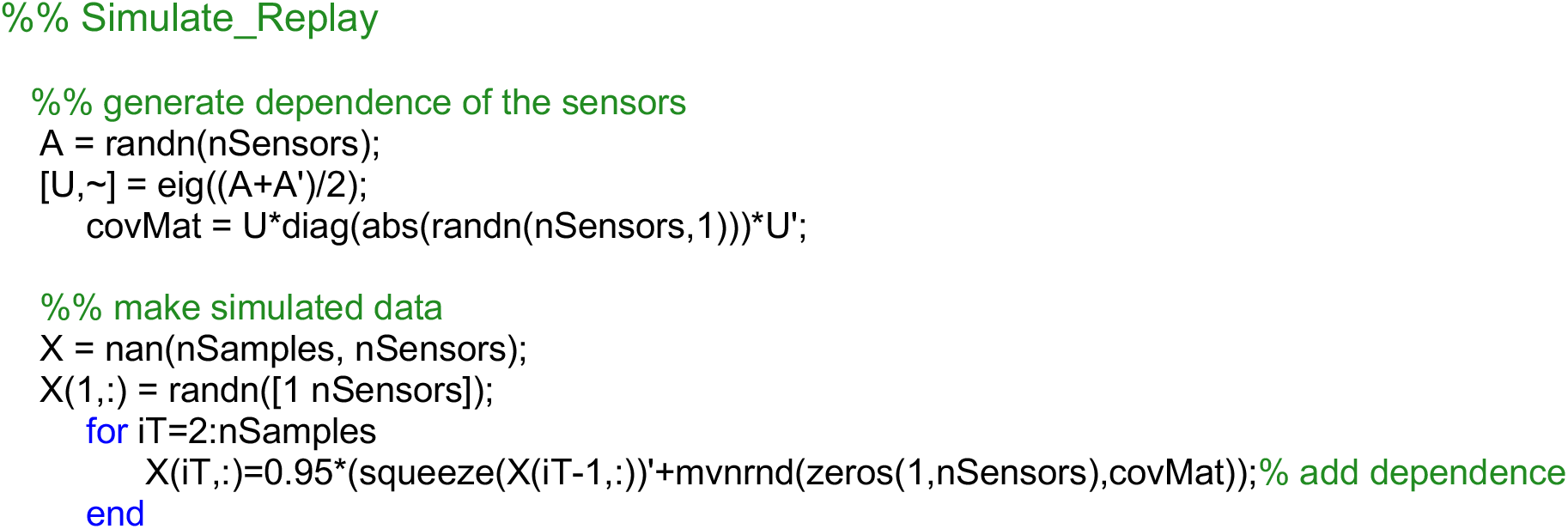

We generate ground truth multivariate patterns of states with core common patterns across all states. This is to respect the fact that most of the states are likely to be defined as pictures, which elicit similar neural activity in general, to some degree. The sequences of the multivariate patterns of states are then injected into the simulated data following the ground truth of state transitions. The state-to-state time lag is assumed to follow gamma distribution with Matlab function “gamrnd”.

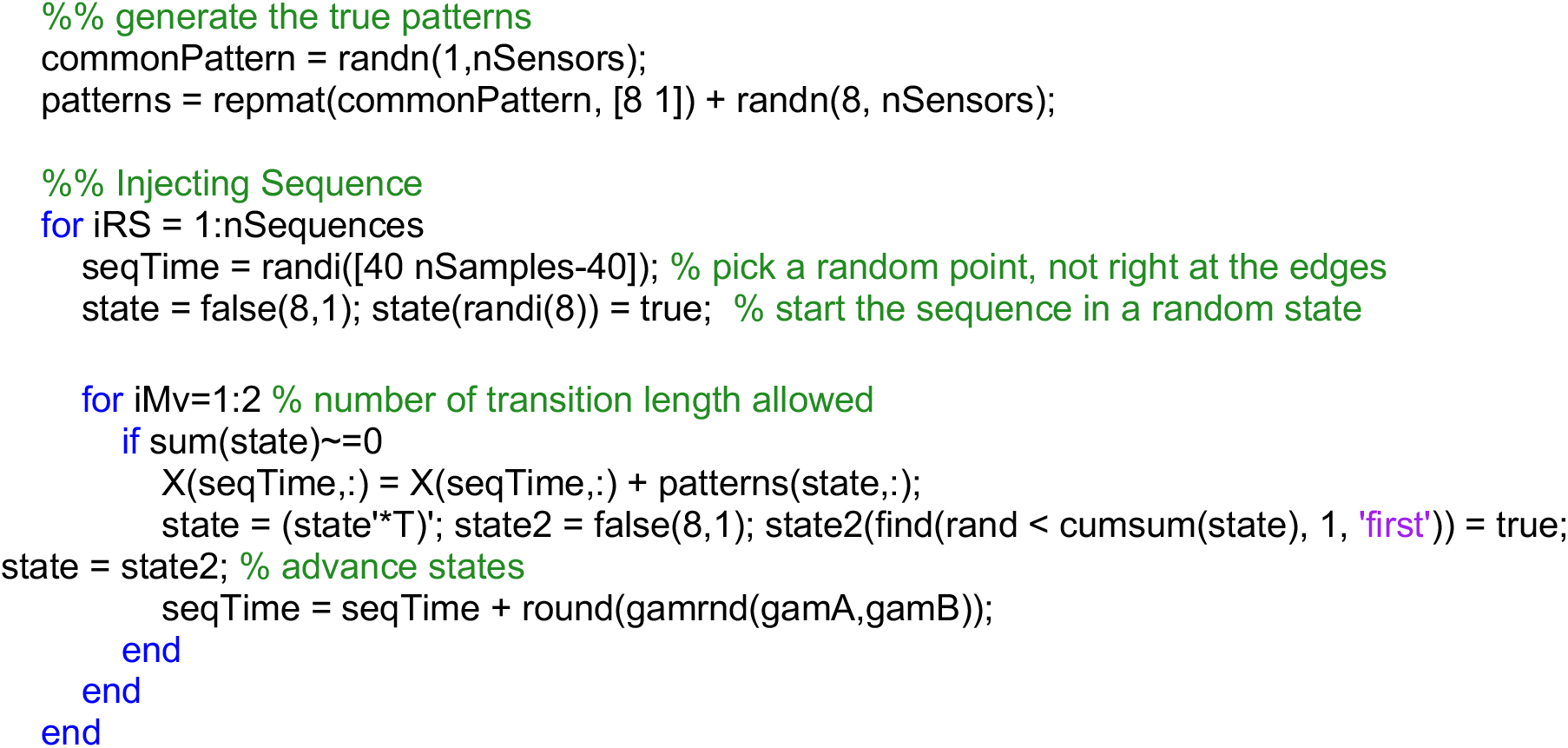

The classifiers are trained based on a different set of simulated data, same as the real MEG data analysis setup. The classifiers are trained using Matlab function “lassoglm”.

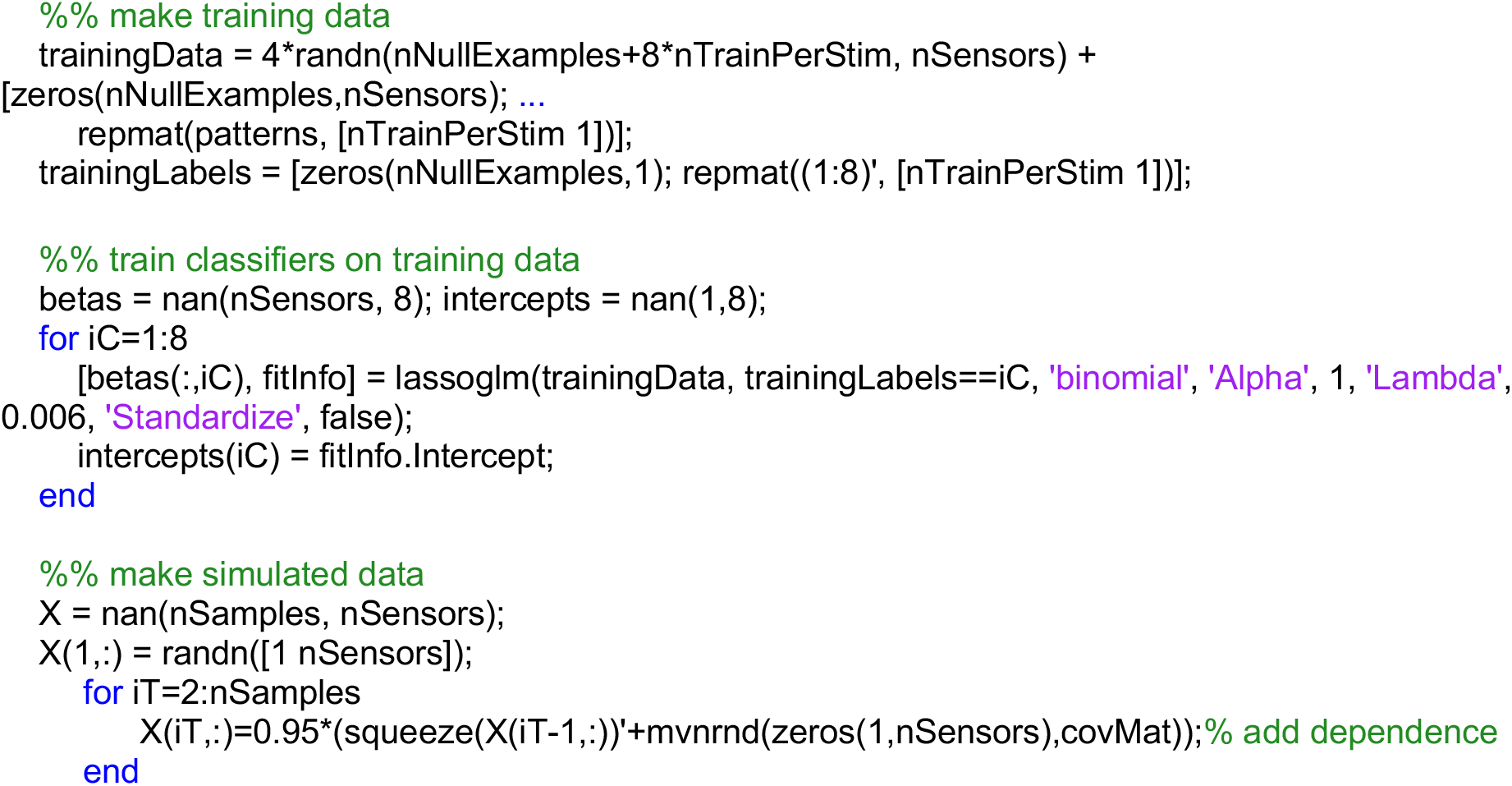

In the end, the simulated data are passed through the trained decoders. The sensor data are transformed to state time courses.

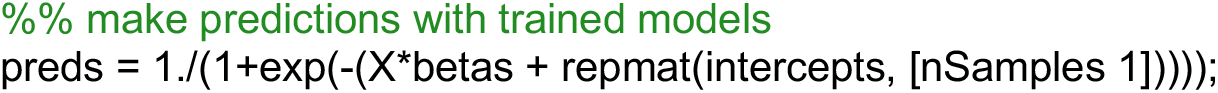

After that, TDLM works on the decoded state space data - \***preds**\* throughout.

## Supplementary Note 2: Rodent data analysis under TDLM

The rodent data is from Matt Wilson’s lab. This is electrophysiology recording from one rat. Data were collected in a spatial navigation task where the rat ran back and forth on a circular track that had a high-wall divider with reward sites on either side. The rat completed 6 rounds of run up (both clockwise and counterclockwise). Fifty-three Cells were recorded in the CA1 of the hippocampus. Spiking activity was recorded at 31,250 Hz /channel. The local field potential was sampled at 2000 Hz. The position of the rat was simultaneously recorded with a sampling rate of 30 Hz. The position records were linearized for later analysis.

The data is analyzed by first subsetting the data based on running speed. In all analysis, the data are restricted to time when the velocity is greater than 10 cm/s. This is to exclude resting or pause during active spatial navigation. We then binned the data for future analysis. The spatial bin size is 5 cm, and the temporal bin size is 10 ms.

The turning curve of the cells on the track is estimated separately based on the running direction (clockwise vs. counterclockwise). After that, the classical one-step Bayesian position decoding is performed on the spike counts of cells, assuming the spike counts follow Poisson distribution, independent between cells and uniform prior over space ^3^. Importantly, the probability of the position is joint estimated based on the running direction and the position on the track. To obtain a readout of the decoded position, we marginalize over the running direction. This is important, because sequence results based on TDLM can be biased by the biased experience (e.g., in clockwise direction, A is always followed by B), estimating jointly with running direction can reduce this concern given the rat has equal experience of running clockwise and counterclockwise.

After that, TDLM can perform on the decoded position space just as the same as on the state time series derived from human MEG data. Importantly, we cannot estimate forward and backward sequence separately, because of the biased experience during decoder training, i.e., the tuning curve at position A is correlated with B, and tuning curve at position B is correlated with C, etc. To correct for this, we rely again on the asymmetry of forward and backward transitions by subtracting reverse sequenceness from forward sequenceness. The state identity permutation test can now work given the rodent running in the clockwise and counterclockwise with equal amount of experience, any asymmetry of forward and backward cannot be explained by the biased experience alone. This analysis pipeline replicates the key rodent finding (Fig 5d), the sequence is forward and is repeating in theta frequency (autocorrelation of sequence peaks at 80 ms, roughly 12 Hz).

## REFERENCES

1 Haxby, J. V., Connolly, A. C. & Guntupalli, J. S. Decoding neural representational spaces using multivariate pattern analysis. Annual review of neuroscience 37, 435–456 (2014).

2 Kriegeskorte, N., Mur, M. & Bandettini, P. A. Representational similarity analysisconnecting the branches of systems neuroscience. Frontiers in systems neuroscience 2, 4 (2008).

3 Biswal, B. B. et al. Toward discovery science of human brain function. Proceedings of the National Academy of Sciences 107, 4734–4739 (2010).

4 Foster, D. J. & Wilson, M. A. Reverse replay of behavioural sequences in hippocampal place cells during the awake state. Nature 440, 680 (2006).

5 Pfeiffer, B. E. & Foster, D. J. Hippocampal place-cell sequences depict future paths to remembered goals. Nature 497, 74 (2013).

6 Skaggs, W. E. & McNaughton, B. L. Replay of neuronal firing sequences in rat hippocampus during sleep following spatial experience. Science 271, 1870–1873 (1996).

7 Behrens, T. E. J. et al. What Is a Cognitive Map? Organizing Knowledge for Flexible Behavior. Neuron 100, 490–509 (2018).

8 Buzsáki, G. & Moser, E. I. Memory, navigation and theta rhythm in the hippocampal-entorhinal system. Nature neuroscience 16, 130 (2013).

9 Carr, M. F., Jadhav, S. P. & Frank, L. M. Hippocampal replay in the awake state: a potential substrate for memory consolidation and retrieval. Nature neuroscience 14, 147 (2011).

10 Ambrose, R. E., Pfeiffer, B. E. & Foster, D. J. Reverse replay of hippocampal place cells is uniquely modulated by changing reward. Neuron 91, 1124–1136 (2016).

11 Kurth-Nelson, Z., Economides, M., Dolan, Raymond J. & Dayan, P. Fast Sequences of Non-spatial State Representations in Humans. Neuron 91, 194–204 (2016).

12 Liu, Y., Dolan, R. J., Kurth-Nelson, Z. & Behrens, T. E. J. Human replay spontaneously reorganizes experience. Cell 178, 640–652 (2019).

13 Mehta, M., Lee, A. & Wilson, M. Role of experience and oscillations in transforming a rate code into a temporal code. Nature 417, 741 (2002).

14 Fyhn, M., Hafting, T., Treves, A., Moser, M.-B. & Moser, E. I. Hippocampal remapping and grid realignment in entorhinal cortex. Nature 446, 190 (2007).

15 Sun, C., Yang, W., Martin, J. & Tonegawa, S. Hippocampal neurons represent events as transferable units of experience. Nature Neuroscience, 1–13 (2020).

16 Colclough, G. L., Brookes, M. J., Smith, S. M. & Woolrich, M. W. A symmetric multivariate leakage correction for MEG connectomes. Neuroimage 117, 439–448 (2015).

17 Deodatis, G. & Shinozuka, M. Auto-regressive model for nonstationary stochastic processes. Journal of engineering mechanics 114, 1995–2012 (1988).

18 Eichler, M. Granger causality and path diagrams for multivariate time series. Journal of Econometrics 137, 334–353 (2007).

19 Lubenov, E. V. & Siapas, A. G. Hippocampal theta oscillations are travelling waves. Nature 459, 534–539 (2009).

20 Wilson, H. R., Blake, R. & Lee, S.-H. Dynamics of travelling waves in visual perception. Nature 412, 907–910 (2001).

21 Weinberger, K. Q., Blitzer, J. & Saul, L. K. in Advances in neural information processing systems. 1473–1480.

22 Vidaurre, D., Smith, S. M. & Woolrich, M. W. Brain network dynamics are hierarchically organized in time. Proceedings of the National Academy of Sciences 114, 12827–12832 (2017).

23 Worsley, K. J. et al. A unified statistical approach for determining significant signals in images of cerebral activation. Human brain mapping 4, 58–73 (1996).

24 Nichols, T. E. Multiple testing corrections, nonparametric methods, and random field theory. Neuroimage 62, 811–815 (2012).

25 Messinger, A., Squire, L. R., Zola, S. M. & Albright, T. D. Neuronal representations of stimulus associations develop in the temporal lobe during learning. Proceedings of the National Academy of Sciences 98, 12239–12244 (2001).

26 Sakai, K. & Miyashita, Y. Neural organization for the long-term memory of paired associates. Nature 354, 152–155 (1991).

27 Barron, H. C., Dolan, R. J. & Behrens, T. E. Online evaluation of novel choices by simultaneous representation of multiple memories. Nature neuroscience 16, 1492 (2013).

28 Kurth-Nelson, Z., Barnes, G., Sejdinovic, D., Dolan, R. & Dayan, P. Temporal structure in associative retrieval. eLife 4, e04919 (2015).

29 Wimmer, G. E. & Shohamy, D. Preference by association: how memory mechanisms in the hippocampus bias decisions. Science 338, 270–273 (2012).

30 Schapiro, A. C., Rogers, T. T., Cordova, N. I., Turk-Browne, N. B. & Botvinick, M. M. Neural representations of events arise from temporal community structure. Nature neuroscience 16, 486 (2013).

31 Garvert, M. M., Dolan, R. J. & Behrens, T. E. A map of abstract relational knowledge in the human hippocampal-entorhinal cortex. Elife 6, e17086 (2017).

32 Penny, W. D., Zeidman, P. & Burgess, N. Forward and backward inference in spatial cognition. PLoS computational biology 9 (2013).

33 McNaughton, B. L., Battaglia, F. P., Jensen, O., Moser, E. I. & Moser, M.-B. Path integration and the neural basis of the’cognitive map’. Nature Reviews Neuroscience 7, 663 (2006).

34 Sirota, A. et al. Entrainment of neocortical neurons and gamma oscillations by the hippocampal theta rhythm. Neuron 60, 683–697 (2008).

35 Buzsáki, G. & Vanderwolf, C. H. Cellular bases of hippocampal EEG in the behaving rat. Brain Research Reviews 6, 139–171 (1983).

36 Baker, A. P. et al. Fast transient networks in spontaneous human brain activity. Elife 3, e01867 (2014).

37 Raichle, M. E. et al. A default mode of brain function. Proceedings of the National Academy of Sciences 98, 676–682 (2001).

38 Tambini, A. & Davachi, L. Awake Reactivation of Prior Experiences Consolidates Memories and Biases Cognition. Trends in cognitive sciences (2019).

39 Norman, K. A., Polyn, S. M., Detre, G. J. & Haxby, J. V. Beyond mind-reading: multivoxel pattern analysis of fMRI data. Trends in cognitive sciences 10, 424–430 (2006).

40 Lewis, P. A. & Durrant, S. J. Overlapping memory replay during sleep builds cognitive schemata. Trends in cognitive sciences 15, 343–351 (2011).

41 Schuck, N. W., Cai, M. B., Wilson, R. C. & Niv, Y. Human orbitofrontal cortex represents a cognitive map of state space. Neuron 91, 1402–1412 (2016).

42 Eldar, E., Bae, G. J., Kurth-Nelson, Z., Dayan, P. & Dolan, R. J. Magnetoencephalography decoding reveals structural differences within integrative decision processes. Nature human behaviour 2, 670–681 (2018).

43 Wimmer, G. E., Liu, Y., Vehar, N., Behrens, T. E. & Dolan, R. J. Episodic memory retrieval is supported by rapid replay of episode content. bioRxiv, 758185 (2019).

44 Foster, D. J. Replay comes of age. Annual Review of Neuroscience 40, 581–602 (2017).

45 Dayan, P. & Daw, N. D. Decision theory, reinforcement learning, and the brain. Cognitive, Affective, & Behavioral Neuroscience 8, 429–453 (2008).

46 Eldar, E., Lièvre, G., Dayan, P. & Dolan, R. J. The roles of online and offline replay in planning. BioRxiv (2020).

47 Schuck, N. W. & Niv, Y. Sequential replay of nonspatial task states in the human hippocampus. Science 364, eaaw5181 (2019).

48 Wittkuhn, L. & Schuck, N. W. Faster than thought: Detecting sub-second activation sequences with sequential fMRI pattern analysis. bioRxiv (2020).

49 Kobak, D. et al. Demixed principal component analysis of neural population data. Elife 5, e10989 (2016).

50 Van Veen, B. D., Van Drongelen, W., Yuchtman, M. & Suzuki, A. Localization of brain electrical activity via linearly constrained minimum variance spatial filtering. IEEE Transactions on biomedical engineering 44, 867–880 (1997).

## Supplementary References

1 Liu, Y., Dolan, R. J., Kurth-Nelson, Z. & Behrens, T. E. J. Human replay spontaneously reorganizes experience. Cell 178, 640–652 (2019).

2 Wimmer, G. E., Liu, Y., Vehar, N., Behrens, T. E. & Dolan, R. J. Episodic memory retrieval is supported by rapid replay of episode content. bioRxiv, 758185 (2019).

3 Zhang, K., Ginzburg, I., McNaughton, B. L. & Sejnowski, T. J. Interpreting neuronal population activity by reconstruction: unified framework with application to hippocampal place cells. Journal of neurophysiology 79, 1017–1044 (1998).

